# A Deep Learning-Driven Sampling Technique to Explore the Phase Space of an RNA Stem-Loop

**DOI:** 10.1101/2024.04.05.588303

**Authors:** Ayush Gupta, Heng Ma, Arvind Ramanathan, Gül H. Zerze

## Abstract

The folding and unfolding of RNA stem-loops are critical biological processes; however, their computational studies are often hampered by the ruggedness of their folding landscape, necessitating long simulation times at the atomistic scale. Here, we adapted DeepDriveMD (DDMD), an advanced deep learning-driven sampling technique originally developed for protein folding, to address the challenges of RNA stem-loop folding. Although tempering- and order parameter-based techniques are commonly used for similar rare event problems, the computational costs and/or the need for *a priori* knowledge about the system often present a challenge in their effective use. DDMD overcomes these challenges by adaptively learning from an ensemble of running MD simulations using generic contact maps as the raw input. DeepDriveMD enables on-the-fly learning of a low-dimensional latent representation and guides the simulation toward the undersampled regions while optimizing the resources to explore the relevant parts of the phase space. We showed that DDMD estimates the free energy landscape of the RNA stem-loop reasonably well at room temperature. Our simulation framework runs at a constant temperature without external biasing potential, hence preserving the information of transition rates, with a computational cost much lower than that of the simulations performed with external biasing potentials. We also introduced a reweighting strategy for obtaining unbiased free energy surfaces and presented a qualitative analysis of the latent space. This analysis showed that the latent space captures the relevant slow degrees of freedom for the RNA folding problem of interest. Finally, throughout the manuscript, we outlined how different parameters are selected and optimized to adapt DDMD for this system. We believe this compendium of decision-making processes will help new users adapt this technique for the rare-event sampling problems of their interest.

## Introduction

Atomistic molecular dynamics (MD) simulations are widely used to study various biological phenomena on the length and timescales often inaccessible by experiments.^1–7^ Some of those phenomena happen on the order of milliseconds. While the limits of conventional bruteforce MD simulations can be pushed to milliseconds with the use of elegantly designed hardware specialized in MD,^1^ conventional MD simulations getting stuck in metastable states still presents a major challenge for users who may not have access to specialized hardware. The high barriers in such free energy surfaces make these transitions rare events that often need advanced sampling methods to accelerate simulations of such systems.^8–12^ Tempering-based^13,14^ or order parameter-based^12,15–17^ sampling techniques are typically used for rare event sampling. While tempering-based sampling techniques do not require prior system knowledge, they cannot overcome large free energy barriers. On the other hand, order parameter-based sampling techniques can easily overcome large barriers if the knowledge of all slow degrees of freedom is made available to the algorithm. One such method is called metadynamics,^12^ which requires defining a set of reaction coordinates or collective variables (CV), i.e., a low-dimensional variable that can distinguish different metastable states of the system.^18,19^ After defining an appropriate set of CV(s) that can capture the slowest degrees of freedom of the system, an external biasing potential that is a function of the CVs is added to the Hamiltonian of the system to push it out of energy well, enabling the simulations to explore other metastable states. Identifying the optimum set of order parameters/CVs for most rare event sampling problems is an active area of research.^20–22^ For that, one often needs to have an *a priori* physical understanding of the process, which may not always be immediately available. To eliminate this need, some studies have used linear dimensionality reduction methods like PCA, TICA, etc. to the simulation data, which generate low dimensional linear embeddings of the system that can be used as a CV. However, such CVs might not be able to capture all the relevant slow degrees of freedom.^23–26^

Non-linear dimensionality reduction techniques like diffusion maps, t-SNE, neural networks, and such are more widely used to preserve the complex couplings in the high-dimensional simulation data, thereby generating a set of CVs that can capture the slowest modes of the process in study.^26–30^ However, the current approaches require the availability of a large training data set from a reference (biased or unbiased) trajectory to determine the CVs. Accordingly, we adapted a novel technique that circumvents this problem. Our approach, DeepDriveMD (DDMD), circumvents this problem by running an ensemble of simulations adaptively, i.e., iteratively running a set of short parallel simulations and seeding new trajectories from undersampled states, as examples applied in different contexts are shown in literature.^31–34^

DDMD is a deep learning-driven MD simulation technique that avoids the problems outlined above by finding undersampled states from an ensemble of running simulations without requiring the user to define a set of physical CV(s).^35,36^ It does so by learning a low-dimensional latent representation^37–39^ of the simulation data on the fly and treating it as a proxy for the CV. DDMD has recently been shown to enhance the conformational sampling of proteins and protein-ligand complexation. ^35^ In this paper, we adapted this method for studying RNA folding, which is a problem fundamentally different than protein folding as RNA folding free energy landscapes are quite rugged compared to that of a protein.^40,41^

RNA plays numerous catalytic roles *in vivo*, which is often believed to be regulated by its folding.^42,43^ The *in vitro* experiments studying RNA folding have reported the presence of non-native low-free energy structures in the folding pathways.^40,44,45^ To get further knowledge about these conformers, atomistic MD simulations prove to be very helpful, as they can provide atomic-level details.^46–54^ The simulations can further provide folding free energy surfaces and folding kinetics, which can facilitate experimental design to understand the folding of RNA molecules.

Here, we report the implementation and efficiency of our approach in simulating the folding pathways of an RNA stem-loop with the sequence of GGCGAGAGCC, henceforth referred to as GAGA in this text. Our results show that the DDMD framework enables the simulations to explore the unexplored parts of the folding/unfolding phase space compared to brute-force MD. This performance of DDMD in enhancing the sampling depends on the learned latent space. Therefore, we also present the qualitative analysis of the latent space, which shows its ability to differentiate between the sampled states. The periodic shifting of simulation resources to undersampled states naturally introduces a bias in the sampling process, which demands a reweighting procedure to estimate the equilibrium populations of the microstates. Therefore, we also introduced a reweighting technique for DDMD inspired by the weighted ensemble simulation method.^55,56^ Finally, we also explained in detail the trials performed to optimize the various components of the DDMD framework for this particular problem. Overall, this paper introduces significant improvements to the purpose of DDMD as a simulation package (as originally it has been mostly used for reaching target states in the least amount of time), and could also serve as a guide to adapt this method to study other complex biological systems.

## Methods

DDMD achieves sampling of undersampled states by using a convolutional variational autoencoder (CVAE) to represent the visited states in a low-dimensional latent space.^22,37,57^ Outliers from this learned latent space are identified and treated as starting points for new simulations, resulting in exploring undersampled regions of the phase space.

DDMD combines MD simulations, ML training, and inference components, as shown in Figure 1. These stages are executed simultaneously, repeatedly, and continuously, with results from each new round of simulations used to (re)train the ML model that is then used to infer from the ongoing simulations and set up the next round of simulations. The ML and inference stages utilize one GPU each. The purpose of the various stages are:

- *Simulation:* performs the MD simulations.
- *Training:* receives data from the Simulation component, (re)trains its ML model (CVAE) with the latest data, and communicates updated model weights to the Inference component.
- *Inference:* aggregates all the MD simulation data and the latest ML model weights to make decisions as to whether to continue or prune a running simulation from the ensemble and assign the initial configuration(s) for the next round of simulations in the Simulation component.

**Figure 1:**
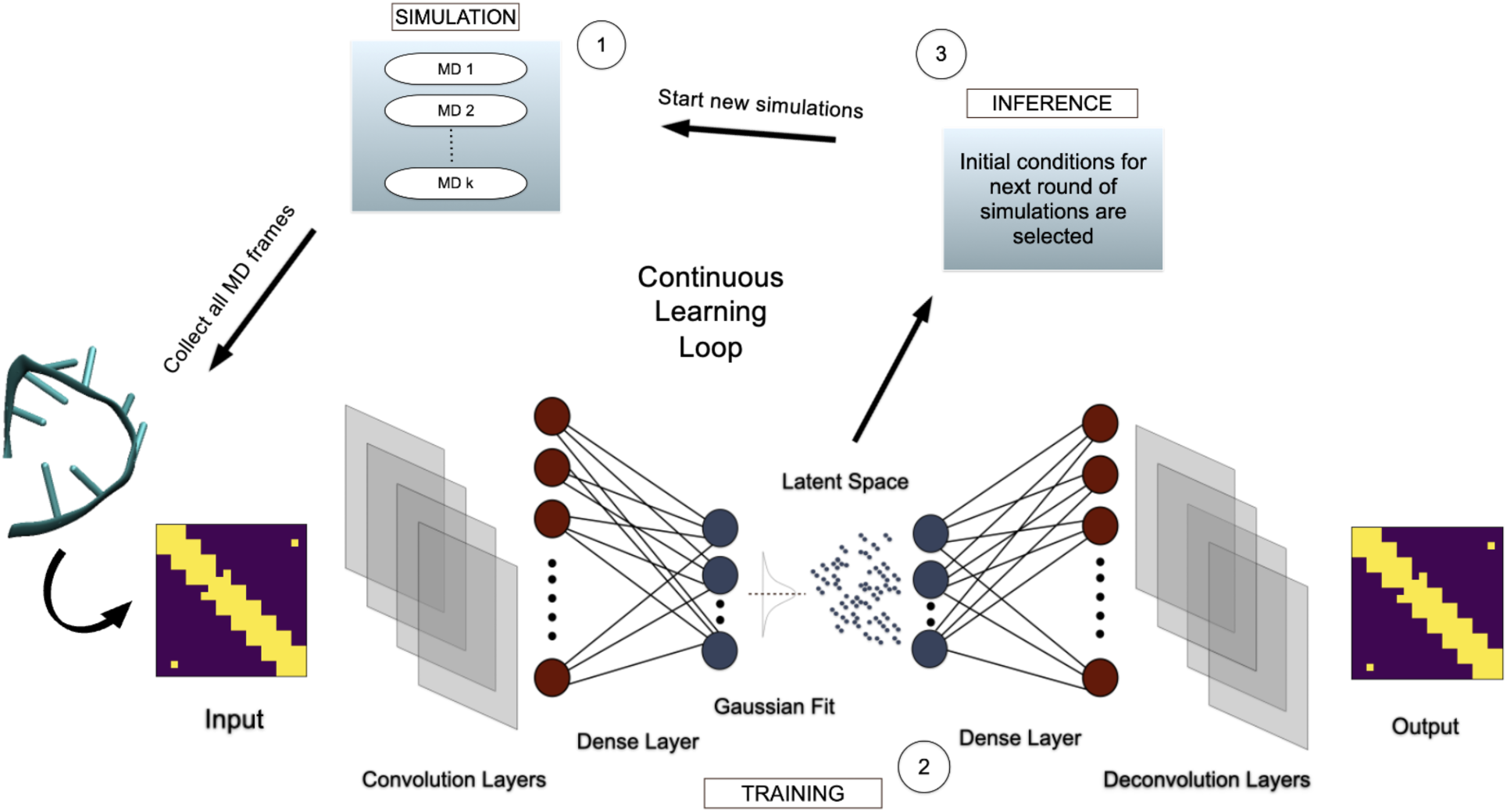
DeepDriveMD framework comprises three stages to support deep learning-coupled simulations.

### MD Implementation

The single-stranded RNA stem-loop studied in this work is a GNRA tetraloop with a GC-based stem, whose complete sequence is GGCGAGAGCC, which will be referred to as GAGA tetraloop hereafter. Similar to the study by Zerze *et al.*,^54^ the Nucleic Acid Builder tool of AmberTools^58^ was used to generate the unfolded initial coordinates of the tetraloop. The sequence was modeled with the nucleic acid force field DESRES and combined with the TIP4PD water model using the library files provided by Kuhrova et al.^59–61^ A single copy of the tetraloop was solvated in a truncated octahedron box of volume 135 nm^3^. Sodium ions were added to maintain electroneutrality, and the salt concentration was adjusted to 1 M of NaCl. The ions were modeled using CHARMM22 parameters.^62^ After solvation and salt addition, the simulation box was equilibrated with 100 ps NVT simulation (T = 300 K) followed by 100 ps NPT simulation (T = 300 K, P = 1 bar). We used OpenMM to perform all the production runs at 300 K and 1 bar.^63^ Electrostatic interactions were calculated using the particle-mesh Ewald method with a real space cutoff distance of 1 nm.^64^ A cutoff distance of 1 nm was also used for the van der Waals interactions. A total of 6 parallel MD simulations were started from the same initial condition, where each simulation runs on a single GPU. This number of concurrently running MD simulations was maintained during the whole course of the DDMD framework. As the simulations progress, the inference part of the algorithm identifies the interesting conformations, i.e., the outliers from the visited conformations, which are potential candidates to start new simulations. Each short simulation was run for 10 ns with a time step of 2 fs.

### ML Implementation

The convolutional variational autoencoder (CVAE) is written in Keras/TensorFlow 2.1.2.^37,65^ It reduces the high dimensional MD trajectory data into a latent vector representation in which structurally similar states cluster together. Among the types of autoencoder, we choose a VAE because it provides several favorable mathematical properties, such as a continuous distribution in the latent space, which are important for statistical analysis of the resulting sampling. In this study, we implemented the CVAE to view every MD frame as a contact map.^66^ Contact maps are built for selected atom pairs. The criteria for atom pair selection for the contact map is presented in the Results section, but the final size of the contact map we used in our production simulations was 20×20. Contacts are considered between each nucleotide (within 7 *Å* cut-off) of the RNA molecule. CVAE learns to represent this contact matrix in a latent space. During the exploration of appropriate CVAE structure, we developed several autoencoder and VAE models, varying the number, width, and type of layers in the model and different sizes of the output latent space, and evaluated them using reconstruction loss. Our final model architecture consists of a symmetric encoder/decoder pair with four convolutional layers. We adjust the convolutional layers based on the size of the system. For our system size, that is, contact map of size 20×20, we use 32 filters with a kernel size of three for all layers and a stride of 1 in all the layers. We then follow the convolutional layers with a single linear layer of 128 neurons and a dropout of 0.5. The CVAE architecture is shown in Figure 1. We systematically tested different latent space dimensionalities ranging from 1 to 10, and the decoder, composed of transposed convolution operators, reconstructs the input contact matrix. We define the loss function as the sum of the binary cross-entropy reconstruction and Kullback-Leibler divergence to an isotropic Gaussian prior N(0, 1). This loss function is optimized using the RMSprop optimizer with a learning rate of 0.001, *⇢* = 0.9, *✏* = 1e-10.

### Inference/ Outlier Detection

The purpose of the Inference component is primarily to select the interesting/outlier conformations to start new simulations. Traditionally, this selection constitutes a biophysical quantity of interest defined as a reaction coordinate, which can be tracked as the simulations run. However, such reaction coordinates are generally system-specific and not always known *a priori* and thus could be challenging to define. To overcome this issue, DDMD uses the CVAE approach to embed the conformations in a low-dimensional manifold starting from a high-dimensional representation defined by generic contact maps. We employed traditional outlier detection methods on the latent embeddings produced by the CVAE to search for undersampled regions of the conformational space. We used the Local Outlier Factor (LOF) algorithm,^67^ as implemented in sci-kit-learn in Python,^68^ to pick the most distant outliers in the latent space. These points represent potentially rare conformations that, if sampled with more MD simulation, could advance the search space. The more negative the LOF is, the higher the outlierness of the particular frame. Hence, we rank all the sampled frames based on their LOF values and select the frames with the most negative LOF values as the starting conditions for the next set of simulations. Computational resources are freed from the trajectories that do not generate any outlier (and hence unproductive), whereas the trajectory that generates an outlier is continued in the next round of simulations. The outlier-generating trajectories are considered productive as they are already sampling a less crowded region in the latent space and, hence, are expected to explore the phase space further if allowed to continue. We note that a particular frame can be identified as an outlier multiple times throughout the simulation.

### Further Filtering of Outliers

Although DDMD does not require prior knowledge of the system, it still offers an option for further refinement of the outliers for systems where some important reaction coordinates are known. For such systems, one can employ any known reaction coordinate(s) to reach a target state, thus combining the data-driven choice with expert knowledge.^69^ For example, one can choose the heavy atom root mean squared deviation (RMSD) to the native state of the RNA as a reaction coordinate to track the progress of simulation on the fly and guide it towards the folded/unfolded state, where the native state represents a structure, determined experimentally via either X-ray crystallography or nuclear magnetic resonance (NMR), against which simulation progress is usually measured. Our primary goal in this work is to get the maximum exploration of the phase space (rather than reaching a target state), assuming no prior knowledge of the system. Accordingly, we presented the free energy surface results of the unrefined cases in the main text. However, we still tested the RMSD-based refining of the outliers (exploitation) combined with outlier-based exploration and presented the results in the Supporting Information (SI).

### Reweighting

By starting short trajectories from new initial conditions and discontinuing ongoing trajectories, the inference component resamples the probability distribution of the whole ensemble of conformations. In doing so, it introduces a statistical bias into the sampling process. To maintain the appropriate Boltzmann distribution of microstate probabilities, we reweight every short trajectory in a similar way as practiced in the weighted ensemble method^34,55,56^ after the sampling is completed. A continuous trajectory, *ζ*, can be considered as a set of discrete time frames as *ζ* = (*x*_0_*, x*_1_*, x*_2_, …., *x*_*n*-1_), where *x_i_* represents the microstate at time *t_i_*, and *n* is the number of frames.^34^ At time t=0 (*t*_0_), the weight (*w*_0_) of the first frame (*x*_0_) of each independent trajectory (we have 6 total) is assigned an equal, fixed, and finite value, *w*_0_ = *c*. We chose *c* = 1, but it is an arbitrary number. All the successive frames will inherit *w*_0_ until the inference component starts working and new trajectories are started from the frames identified as outliers. Then the reweighting proceeds as follows. Let’s consider a short trajectory *ζ*′ with *n* frames (*n* is the same for all the short trajectories). The *j*^th^ frame of *ζ*′ is identified as an outlier and *N* new trajectories are started (each of these new trajectories will emerge at a different point of time in the whole simulation) by taking that outlier frame as the starting configuration. The weights of the frames of *ζ*′ numbered from *j* +1 to *n* are updated to *w_j_/*(*N* + 1), where *w_j_* represents the weight of the *j*^th^ frame. The weight of each of the trajectories which starts from that particular *j*^th^ frame of *ζ*′ is also set to *w_j_/*(*N* + 1) and so on. In this manner, the new weights are assigned to each frame, conserving the appropriate statistical distribution of states in all the trajectories. The reweighting scheme is shown in Figure 2.

**Figure 2:**
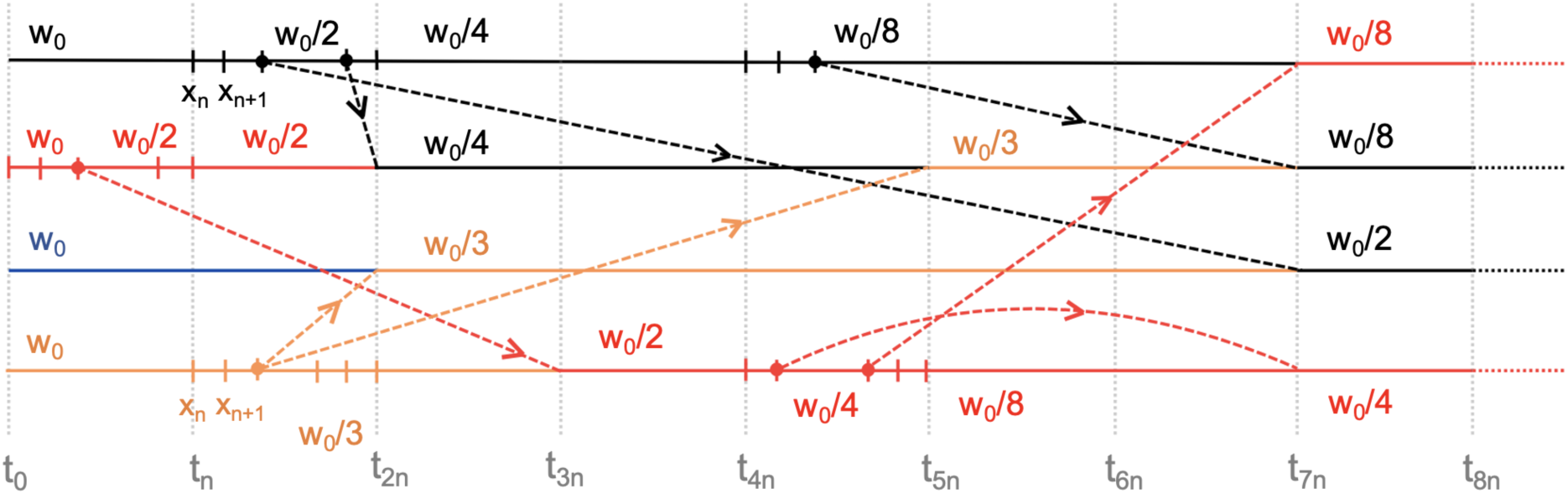
Reweighting example for 4 walkers: The initial 4 colors (black, red, blue, and orange) represent the different walkers which started from the respective initial conditions. The dashed arrows represent the source of the outlier, and their ends show the beginning of the respective new trajectories. The weights *w_i_* represent the weights of the respective parts of the trajectories. The gray vertical dotted lines represent the starting time of every short trajectory (of n frames).

### Analysis Methods

We quantified the free energy surfaces (FES) of the tetraloop using two order parameters, one of which is a similarity-based order parameter (Q), and the other one is the heavy atom root-mean-square distance (RMSD) from the reference native structure. The first of the 10 NMR structures present in the PDB entry 1ZIG was used as the reference folded structure for the GAGA tetraloop.^70^ Q only encompasses knowledge that the stem folds into a canonical A-RNA configuration. We defined Q only for the stem part of the tetraloop following a protocol analogous to the related study done by Zerze et al.^54^ The order parameter Q is defined as

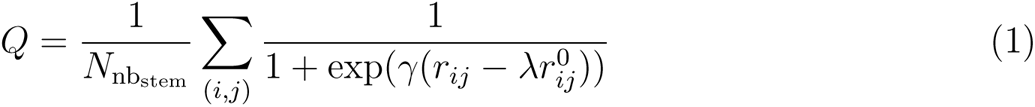

The generalized definition of Q is adapted here for the interstrand contacts.^71,72^ The sum runs over *N_nbstem_* pairs, which is the total number of (stem) atomic pairs (*i, j*) that are considered in contact in the canonical A-RNA state. Any heavy (i.e. non-hydrogen) nucleobase (nb) atom of the 3’ strand was considered in contact with a heavy nb atom of the 5’ strand if the distance between them is less than 5 Å in the reference native structure. *r^0^ij* and *r_ij_* are the distances between *i* and *j* in the reference native structure and in any given instantaneous configuration, respectively. *r* in the smoothing function was taken as 50 nm^-1^ and the adjustable parameter *A* was taken as 1.5.^72^

The unbiased two-dimensional probability densities of order parameters Q and RMSD were obtained after reweighting all the trajectories as mentioned in the above section. The unbiased probability density, P(Q, RMSD), was then converted to free energy via the equation

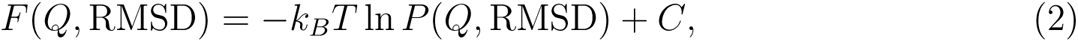

where *k_B_T* is the product of the Boltzmann constant and the temperature, and *C* is an immaterial constant. Since *Q* only includes the stem atoms, RMSD is particularly useful in quantifying the structural differences arising from the loop part, as this feature includes both stem and loop atoms. The reference paper defines the relevant states for this system based on these two order parameters.^54^ Accordingly, we consider the misfolded state (M) to have Q greater than 0.9 and RMSD between 0.3-0.4 nm, while the folded state (F) has Q greater than 0.9 and RMSD less than 0.2 nm.

## Results and Discussion

### Atom Selection for Contact Maps

We first tested various atom selection criteria for the contact map to train the CVAE. To determine the best atom pairs, we used the lowest RMSD achieved by the workflow in a fixed wall clock time (4 days in this case) as the criterion, where an unfolded state (Q *⇠* 0, i.e., a high RMSD configuration) was the initial condition (IC). The idea behind this approach is that a better atom selection will lead to a better-trained CVAE and hence, will better represent an MD frame in the latent space. A better-represented latent space will enable the inference stage to generate outliers, guiding the system to sample undersampled regions of the phase space and enabling the escape from local free energy basins. Consequently, the system should be able to sample all the relevant states leading to the farthest configuration. The atom selection criterion enabling the workflow to sample the minimum value of RMSD during its runtime was chosen as the most appropriate. Table 1 shows the different atom selections tested with their corresponding lowest RMSD sampled by the workflow. The atom pair C1’ and C3’ was observed to sample the least value of the lowest RMSD value in a wall-clock runtime of 4 days and, therefore, was selected to run production simulations.

**Table 1:** Lowest RMSD to the native state sampled for different atom selections. The atom combination C1’ and C3’ was chosen as the best one as it enabled the framework to get closest to the folded state (i.e., farthest to the starting condition) in a fixed wall clock time.

| Atom Selection | Lowest RMSD (nm) |
| --- | --- |
| C2' and O5' | 0.3 |
| C1' and O5' | 0.38 |
| C3' and N1 | 0.5 |
| O5' and N1 | 0.52 |
| C1' and P | 0.56 |
| C1' and C3' | 0.27 |

### Dimensionality of the Latent Space

To determine the optimal latent space dimensionality for a reasonable representation of a contact map by the CVAE, we tested the effect of changing the latent space dimension on the overall loss (i.e. the sum of reconstruction loss and the latent loss) of the CVAE.^37^ Using cross-entropy loss, the reconstruction loss is used to gauge the ability of the CVAE to reconstruct the original input contact matrix. The latent loss accounts for the loss in information when the latent space embeddings are forced to fit a Gaussian distribution. It is calculated as the Kullback-Leibler (KL) divergence between the latent embeddings and a normal distribution with a mean of 0 and a standard deviation of 1. We used a 5 *µ*s brute force MD trajectory of GAGA tetraloop to train the CVAE and calculate its overall loss in creating a latent space of dimensionalities ranging from 1 to 12. As expected, the loss decreased on increasing the latent space dimensionality as shown in Figure 3. This presents a trade-off between the compressibility of the MD simulation data and the loss of information after compression because although a higher number of latent space dimensions will better present the data, it will be computationally more expensive to train the CVAE as the latent space dimensionality increases. Additionally, a higher latent space dimensionality than needed will overcomplicate the analysis of the latent space. Therefore, we chose the optimal latent dimensionality as 6.

**Figure 3:**
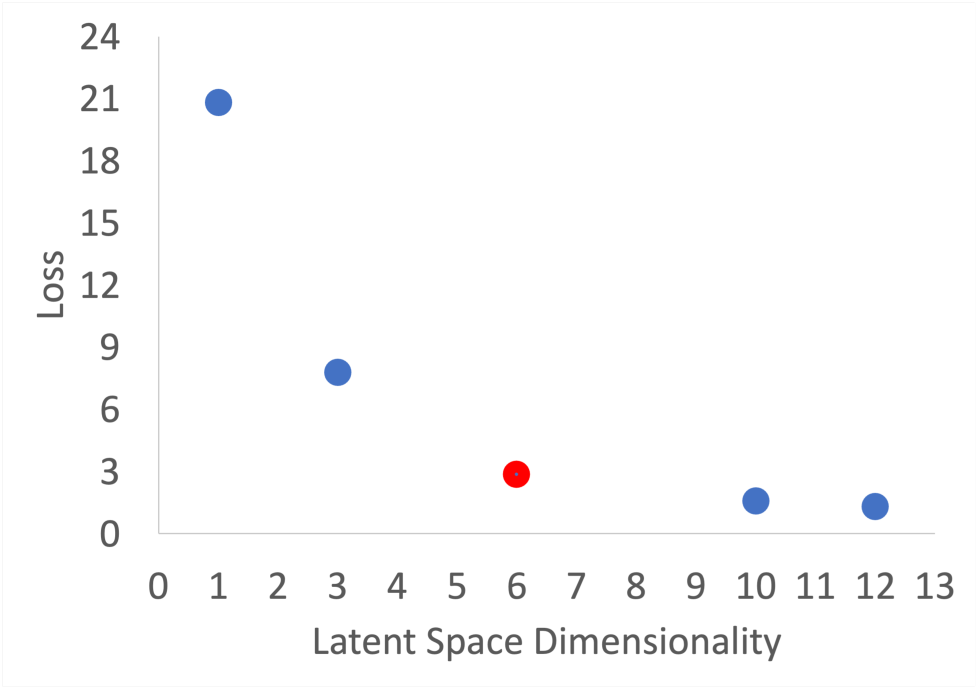
Mean reconstruction loss at various latent dimensionalities showing the trade-off between compression and model accuracy. The higher the latent space dimensionality, the better the representation of the contact maps but training the CVAE becomes computationally more expensive. We chose the latent dimensionality of 6, after which the loss decrease was minimal.

### Free Energy Surface of Folding and Unfolding

We performed multiple sets (7 total) of simulations with various initial conditions (IC) in each set to reproduce the free energy surface of the GAGA tetraloop. We labeled the sets of simulations as sx (x being an integer between 1 and 7). Set s5 had 3 of the walkers’ ICs having Q *⇠* 0.5, while the other 3 had ICs of a folded configuration (Q *⇠* 0.99). In set s6, 3 walkers had the IC having Q *⇠* 0.75, while the other 3 walkers started with a folded configuration (Q *⇠* 0.99) as the IC. Each walker in sets s5 and s6 was simulated for 1.7 and 1.4 *µ*s, respectively. To argue if DDMD can faithfully sample the whole phase space, we quantitatively compared the FES obtained from each DDMD simulation to that obtained using Parallel tempering in the well-tempered ensemble combined with the well-tempered metadynamics (PTWTE-WTM)^12–15,73,74^ in the study by Zerze *et al.*^54^ For this comparison, we calculated the absolute free energy difference (FED) between the DDMD and PTWTE-WTM simulations for each grid point.^75^ The reweighted FES for sets s5 and s6 and the corresponding FED are shown in Figure 4. The FES of GAGA tetraloop has folded (F), unfolded (U), and misfolded (M) basins as defined in our previous work.^54^ These FESs can be seen to capture all the relevant free energy basins involved in the folding/unfolding landscape of GAGA tetraloop. From the FEDs, we observe that the free energies of the U, M, and F regions are captured well, which are the relevant states for this system. The FEDs show that the differences in transition state free energies are relatively large.

**Figure 4:**
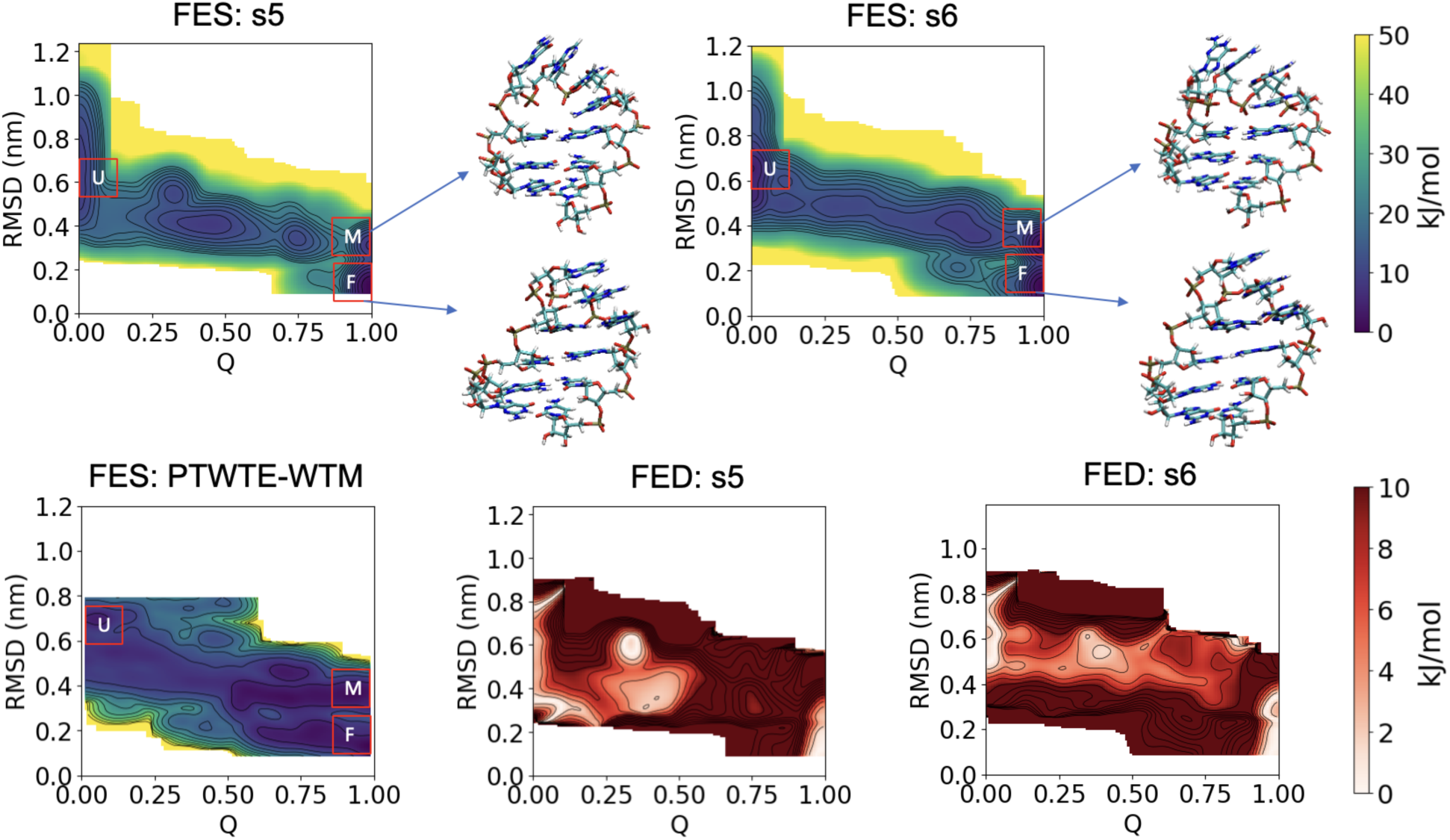
The FES (top panel) for sets s5 and s6, along with the representative configurations of the M and F states; and the corresponding absolute free-energy difference (FED) (bottom panel). The reference PTWTE-WTM FES^54^ is shown in the bottom left. We capped the FED at 10 kJ/mol to aid the visualization. The FES obtained from DDMD captures all the relevant free energy basins involved in the folding/unfolding landscape of GAGA tetraloop.

We also calculated the FE difference between folded and unfolded states, that is, Δ*F_FU_*, using the 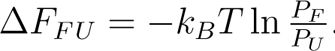.^75^ Here, *P_F_* and *P_U_* are the unbiased probabilities of finding the system in these states at the simulated temperature. Δ*F_FU_* was found to be -6.4 kJ/mol in the reference PTWTE-WTM simulation, while it is -6.8 kJ/mol and -3 kJ/mol for simulations 5 and 6, respectively, which also shows a reasonable agreement between DDMD and PTWTE-WTM.

While these findings are highly promising (especially given the reduced costs as discussed in *Computational Cost* subsection), we noted some initial condition dependence of the findings. RNA folding transition is notorious for being glass-like, and the initial condition dependence is not an uncommon problem for some other sampling strategies.^40,45,54^ We further analyzed the extent of initial condition dependence from additional simulations (sets s1 to s4), where we restricted the ICs of each walker to a particular configuration. Specifically, we used four separate ICs corresponding to Q *⇠* 0, 0.5, 0.75, and 0.99 for s1, s2, s3, and s4, respectively. ICs of each walker within a given set were kept identical for these sets. FES presented in Figure S1 indicated that using these sets did not reproduce the FES of the GAGA tetraloop. Therefore, we concluded that variation in the ICs is essential for properly reproducing the FES. We note that sampling through DDMD is driven by the selection of outliers, which is limited by the variability in the latent space. One of the limitations of the current version of the DDMD code is its inability to adaptively “forget” portions of the conformational space that constitute productive sampling pathways.^35^ Consequently, this limitation results in less well-explored FESs or IC-dependence for relatively more complicated glacial systems like the RNA in this work. We are working towards including adaptive memory models to improve this within future versions of DDMD.

To demonstrate the power of DDMD in enabling the simulation to sample a target configuration (i.e. the folded state), we also performed an outlier-driven exploration simulation combined with RMSD refining of the outliers. For this, an unfolded state (Q *⇠* 0, i.e., a high RMSD configuration) was taken as the IC. In every iteration of the inference stage, the top outliers are further arranged in the order of their heavy atom RMSD to the native state to enable the workflow to prioritize the lower RMSD frames as the initial conditions of the next round of simulations. This approach enables the system to sample a configuration having RMSD as low as 0.1 nm in a wall-clock time of 6 days (1.9 *µ*s per walker). This is a significantly improved sampling speed compared to a brute-force simulation. The corresponding FES is shown in Figure S2. The outliers filtering pushes the system to reach a target state at the cost of lowered exploration ability of the workflow. Nonetheless, the FES shows some low free energy basins, which correspond to (Q *⇠* 0.5, 0.3 *<* RMSD *<* 0.4), (Q *⇠* 0.75, 0.3 *<* RMSD *<* 0.4) and (Q *⇠* 0.99, 0.1 *<* RMSD *<* 0.3) (Figure S3).

### DeepDriveMD Enhances the Sampling of Transition States Compared to MD

Although some basins remained unexplored in sets s1 to s4, they still offered significantly better exploration than conventional brute-force MD simulations of the same length and number of replicas (i.e., for the same cost); starting from the same ICs. Figure 5 shows two-dimensional and one-dimensional histograms (Q and heavy atom RMSD) of the conformations sampled by DDMD (s1, s2, and s4) and brute-force MD simulations (starting from the same corresponding ICs with the same number of independent walkers, which is 6, and histograms are calculated from data combining all 6 walkers). For set s1 (IC has Q *⇠* 0 and RMSD *⇠* 0.86 nm), the brute-force simulation keeps sampling in the same region from where it started, as unimodal distributions of the parameters indicate. In contrast, DDMD enables the simulation to sample more diverse conformations and, hence, newer parts of the phase space, as depicted by the multimodal distribution. We had a similar observation for set s4 (IC Q *⇠* 0 and RMSD *<* 0.2 nm), where DDMD enables enhanced sampling of the transition states having higher RMSD compared to brute-force simulation (Figure 5). We had a rather interesting observation for set s2 (IC Q *⇠* 0.5). The brute-force MD simulation, in this case, ends up sampling the energy wells corresponding to the unfolded region and the region having RMSD in the range of 0.25-0.35. The histogram distribution for set 2 shows that DDMD performs well in pushing the system to sample the transitions in the middle RMSD range, too, thereby leading to a much better free-energy estimate of most states (Figure S1).

**Figure 5:**
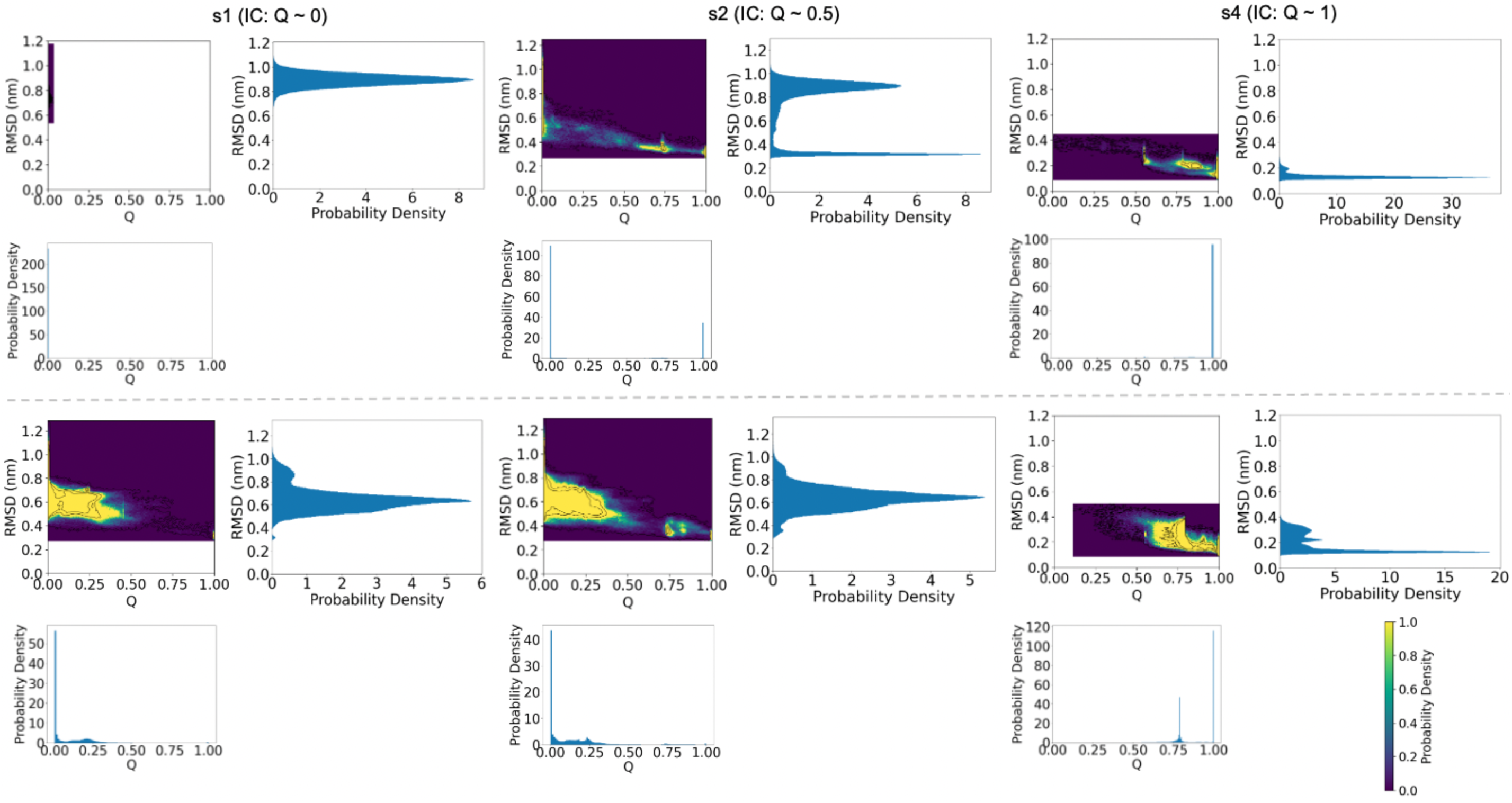
The 2-D probability densities and the corresponding projections on 1-D for brute-force simulations (top panels) and DDMD (bottom panels) for the specified sets of simulations. The color bar shown is for the probability density in the 2-D projections.

### Representation of Conformations in the Latent Space

Sampling through DDMD is driven by the selection of outliers which are highly sensitive to the quality of latent space in which all the sampled conformations are represented. Hence, the latent space must capture the structural similarities/dissimilarities between the conformations. To verify this, we project the latent space embeddings into a 2-dimensional space using the t-distributed stochastic neighborhood embedding (t-SNE) method,^76^ and color this space using the two order parameters, Q and heavy atom RMSD to the native state (Figure 6). The colored latent space representation corresponding to each of the simulations shows a good segregation of conformations with respect to RMSD. The conformations having low, high, and medium values of RMSD can be seen predominantly in specific regions of the projected latent space. The colored latent space representations for sets s1-s4 are shown in Figure S5. The segregation of conformations with respect to Q is less apparent for sets s1, s2, s3, s5, and s6 since they all sample the Q *<* 0.1 region, which is a highly degenerate state, and hence it populates almost the whole latent space. Nevertheless, differently-colored pockets corresponding to low, middle, and high Q conformations can be seen in all projected latent spaces. This observation is crucial as it exhibits the ability of CVAE to represent the RNA conformations in clusters without having any knowledge of the target state. We also note that the projected latent space for s2 and s3 looks very similar in terms of overall shape and location of middle and high Q conformations, which could mean that a trained CVAE model is transferable to another independent set of simulations. The similarity of latent space also supports the fact that the CVAE learns the reaction coordinates pertaining to the folding/unfolding problem with reproducibility.

**Figure 6:**
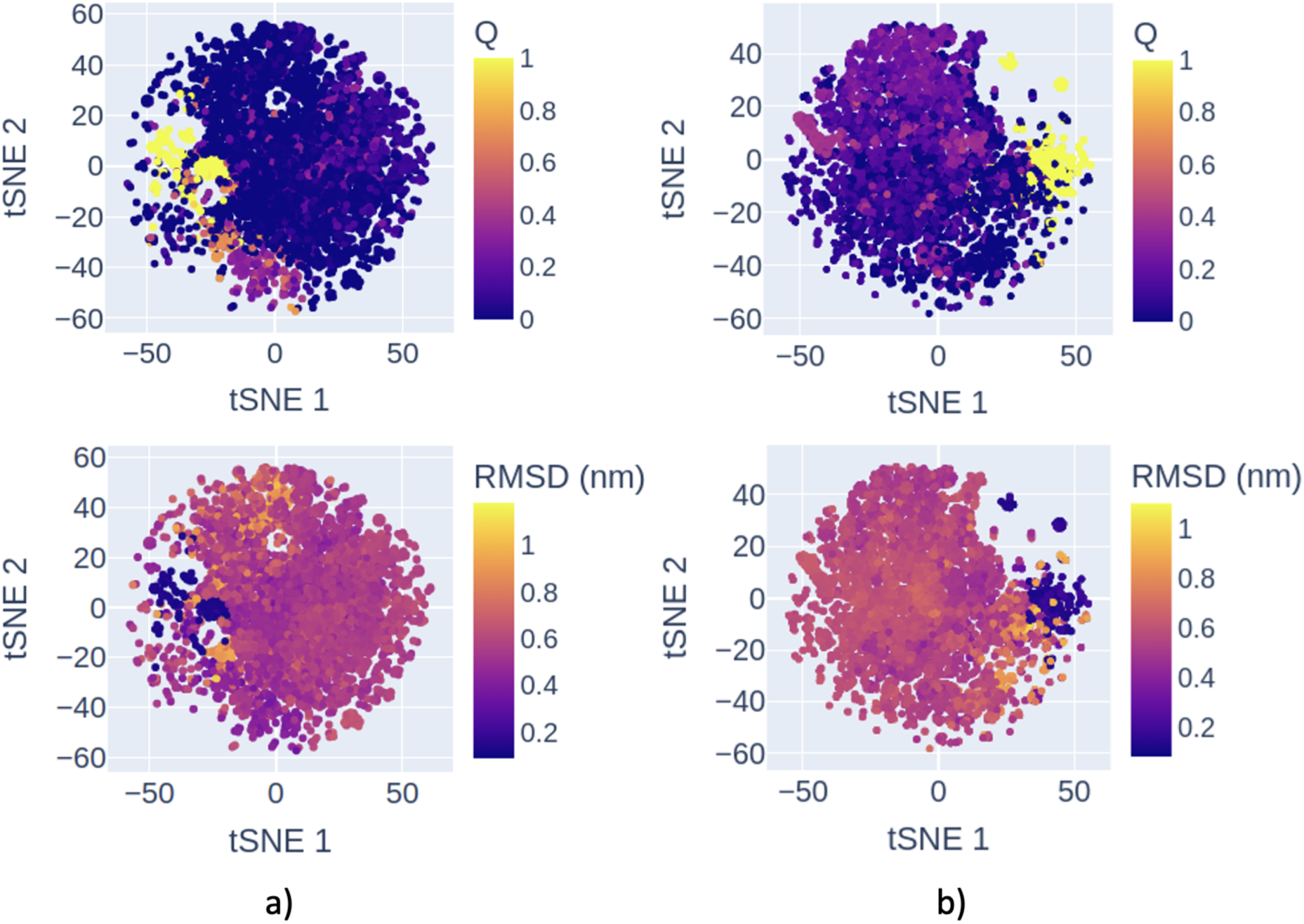
Latent space representations of conformations sampled by a) s5, and b) s6, projected on a 2-D space using tSNE. The latent space has been colored separately using the two order parameters, Q (top panel) and the heavy atom RMSD to the native state (bottom panel). The plots show that the latent space can segregate the conformations based on the respective order parameters.

### Structural Differences between Outliers and Inliers

Having established the quality of the latent space, we sought to verify if the LOF can indeed identify the relevant/interesting conformations out of a fairly large collection of them. In principle, the more structurally dissimilar a conformation is compared to the pool, the higher its ‘outlierness’ should be. To test this, we picked random outliers as identified by our simulations. For each outlier, we selected representative inliers by clustering the remaining pool (i.e. all the conformations sampled before the selection of the particular outlier) using k-means,^77^ and then selected the points closest to the largest 3 clusters as the 3 inliers. We used Barnaba^78^ to characterize the selected conformers. Specifically, we analyzed the base pairs and stacked pairs to find a common structural feature present in the 3 inliers but absent in the outlier or vice versa. For most cases, we found that outliers had at least one structural difference (stacked pairs or/and base pairings) compared to the representative inliers, as shown by the inlier and outlier conformations for sets s1 and s2 in Figure 7. We had similar observations for other sets too. These findings support the ability of LOF to identify structural outliers from a huge pool of RNA conformations.

**Figure 7:**
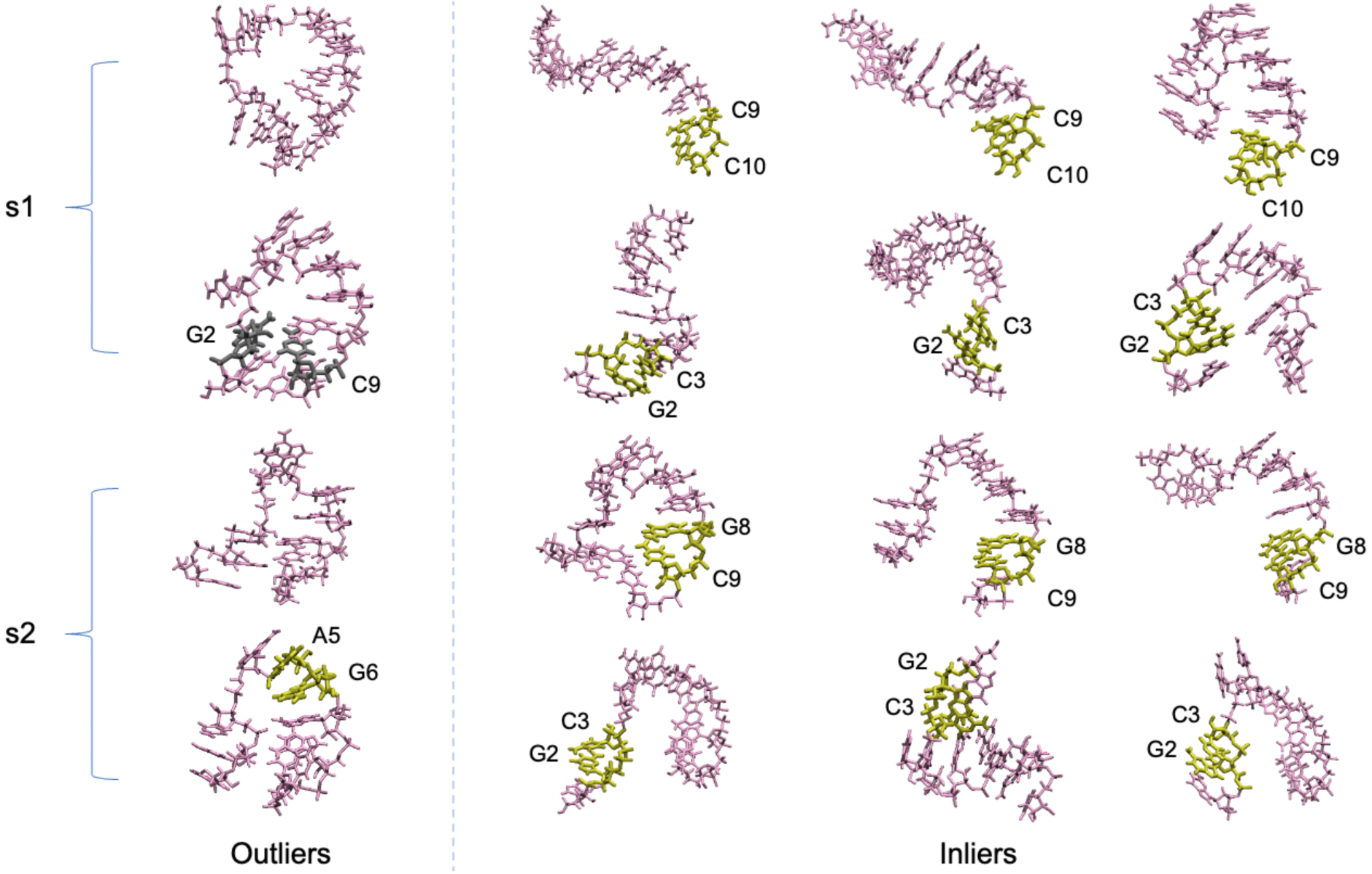
Structural deference’s between outliers and inliers: The outlier conformations are on the left of the dashed line, while the corresponding 3 representative inliers are on the right side of the dashed line. Only the base pairs (gray) and the stacked pairs (yellow), which are present in the outlier but missing in the corresponding 3 inliers and vice-versa, are shown.

### Structural Characterization of the High-Q Regions

As another quality test of the latent space, we sought to characterize the high-Q conformations using their latent space embeddings. Zerze *et al.*^54^ reported that the most populated structure of M has its first 3 and the other 7 nucleobases separately stacked together, while for the F region, the most populated structure has its first 4 and the other 6 nucleobases stacked separately together. To check if the latent space can capture this distinction, we found the most populated structures corresponding to M and F by performing k-means clustering of the latent space embeddings corresponding to those defined regions separately. We considered the embedding closest to the centroid of the biggest cluster to be the representative structures for the M and F energy basins (Figure 4). The representative M and F structures from DDMD simulations here showed the same stacking as in our previous work.^54^ In agreement with the reference study, we also found a lack of the noncanonical A-G base pair in the loop portion of our M structure.

### Computational Cost

We compared the computational cost with which our simulations sampled the FES for GAGA folding/unfolding to two other computational studies. One of them is the main reference paper for the present study, in which the PTWTE-WTM method was used, while the other one employed the MM-OPES simulation method.^16,75^ The MM-OPES method has been reported to have lower computational costs when compared to PTWTE-WTM in terms of CPU and GPU hours: *⇠* 18250 CPU hours and *⇠* 2600 GPU hours used by PTWTE-WTM, while *⇠* 3800 CPU hours and *⇠* 630 GPU hours used by MM-OPES (Table 2). These hours represent the combined wall clock time for which all the CPUs and GPUs were occupied to generate the converged FES. To test the convergence point for our simulations, we calculated our FES’s average free energy error with time by taking the final FES (i.e. obtained after the total simulation time per replica) as the reference (Figure S6). The simulation was considered to converge when the average error settled down to *⇠*0 kJ/mol. From Figure S6, we inferred that the FES of simulations 5 and 6 converged in 0.25 and 0.2 *µ*s, respectively. We calculated the computational cost of generating the converged FES from simulations 5 and 6. Each simulation was run on a node having 48 Intel Xeon G6252 CPUs and 8 Nvidia V100 GPUs. To calculate the CPU/GPU hours, we used the formula

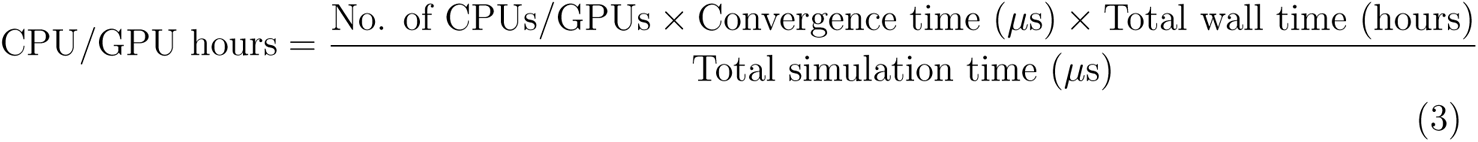

**Table 2:** The computational cost of DDMD compared to PTWTE-WTM and MM-OPES.

| Simulation | CPU Hours | GPU Hours |
| --- | --- | --- |
| PTWTE-WTM | 18250 | 2600 |
| MM-OPES | 3800 | 630 |
| Simulation 5 (DDMD) | 935 | 156 |
| Simulation 6 (DDMD) | 720 | 120 |

For the converged FES of simulations 5 and 6, we reported the CPU and GPU hours in Table 2. Our method is almost 20 times less costly than the PTWTE-WTM method and 4 times less costly than MM-OPES.

## Conclusions

Due to their capacity to enable smarter computational campaigns and thereby accelerate scientific discovery, ML methods play a more prominent and significant part in computational modeling. Such approaches are effective because they provide straightforward, scalable, and broadly applicable ways to handle high-dimensional and potentially high-volume datasets. This ability is crucial when working with biomolecular systems because the simulation datasets used in these studies are high dimensional. In this work, we have demonstrated the ability of DeepDriveMD (DDMD), a general-purpose and extensible framework for implementing ML-driven simulations in sampling the folding pathways of the GAGA tetraloop. Studying folding FES of RNA through atomistic simulations is nontrivial, owing to kinetic traps and relatively deep metastable states in the energy landscape. We have shown that DDMD addresses this issue by pushing the system to sample other transition states while preserving the sampling rigor of the low free energy states. It does that by on-the-fly learning of the low-dimensional latent space from the running simulation itself, without requiring the user to define any *a priori* order parameter. Traditional parallel tempering (temperature replica exchange) simulations rely on extensive use of thermal fluctuations ^13^ and metadynamics variants modify the free energy distribution of the system to push it out of the low free energy states.^12,14,73^ These methods modify the Hamiltonian of the system to maximize the number of visited microstates. The working of DDMD fundamentally differs in this aspect as it facilitates enhanced sampling by only choosing new starting configurations from the already sampled ones (no modifications to the Hamiltonian or no exchanges in the temperature space). This allows a successful sampling scheme only with minimal bias in the sampling process and also preserves the kinetics of the system.

We note that the ability of the autoencoder to learn new information also affects the decision to identify the undersampled states. As the amount of simulation data increases, the memory dedicated to latent space gets saturated, leading to nearly static latent space, which leads to substantially lowered ‘outlierness’ for the selected outliers, hampering the exploration of phase space, which affected our simulations as well (e.g., the initial condition dependence).

Making the CVAE more dynamic in the presence of a large amount of simulation data is one of our future goals that will result in reliable FES with no initial condition dependence.

Our results show that DDMD enables much better sampling of the GAGA tetraloop compared to brute-force MD simulations while having a significantly lower computational cost than the state-of-the-art methods. The algorithm could sample a reasonably low RMSD frame without even requiring any knowledge of the folded state, which would help predict the structures of RNA molecules whose native state is not much researched. Such a goal could be accelerated further if a combination of exploration and exploitation is used to enable the simulations to reach a target state. As an added advantage, DDMD executes the simulations at a constant temperature, and thus the reweighted microstate populations allow the user to study the kinetics involved in the process. Markov state model (MSM) is a valuable tool that can be used to estimate the transition rates of the metastable states using the reweighted equilibrium populations.^79,80^ MSMs have also been used to drive the adaptive sampling of short protein folding simulations, hence reducing the uncertainty of thermodynamic observables.^81^

Throughout this paper, we presented the methodology to extend the use of DDMD (which has until now been used to study protein-folding and ligand binding) to study the fundamentally different problem of RNA folding. We hope this can serve as a useful guide for adapting this method to study bigger RNA molecules and even other types of complex systems, for instance, protein-RNA complexation and biomolecular condensates.

## Acknowledgement

This work at the University of Houston is supported by funding from the Cancer Prevention and Research Institute of Texas (CPRIT) award RR220008 and the Welch Foundation Catalyst Center for Advanced Bioactive Materials Crystallization (Award V-E-0001) to G.H.Z. The simulations presented in this work were performed on the computational resources provided by the Hewlett-Packard Enterprise Data Science Institute at the University of Houston.

## Supporting Information Available

6 Supporting Figures are available.

## TOC Graphic

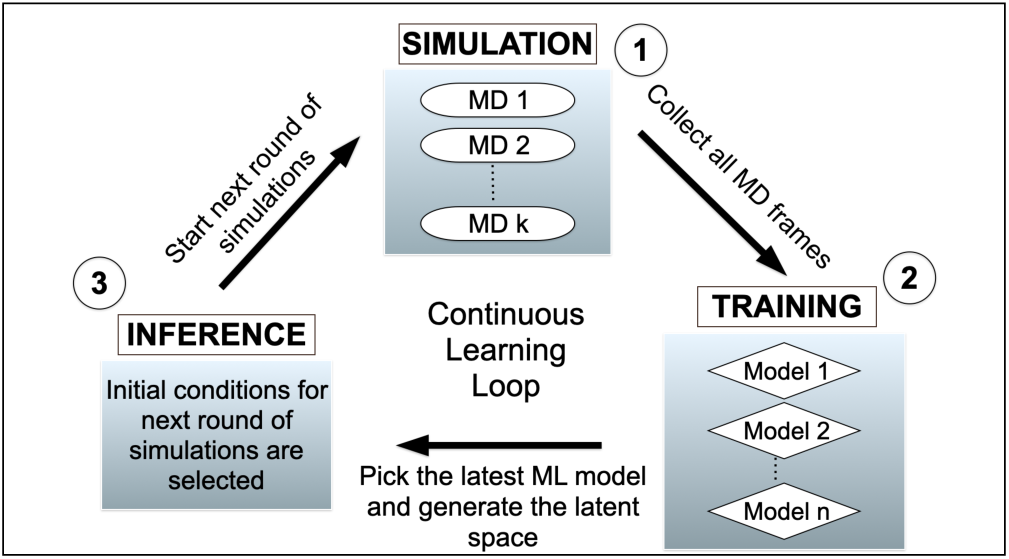

## Supporting information

### Supporting Figures

**Figure S1:**
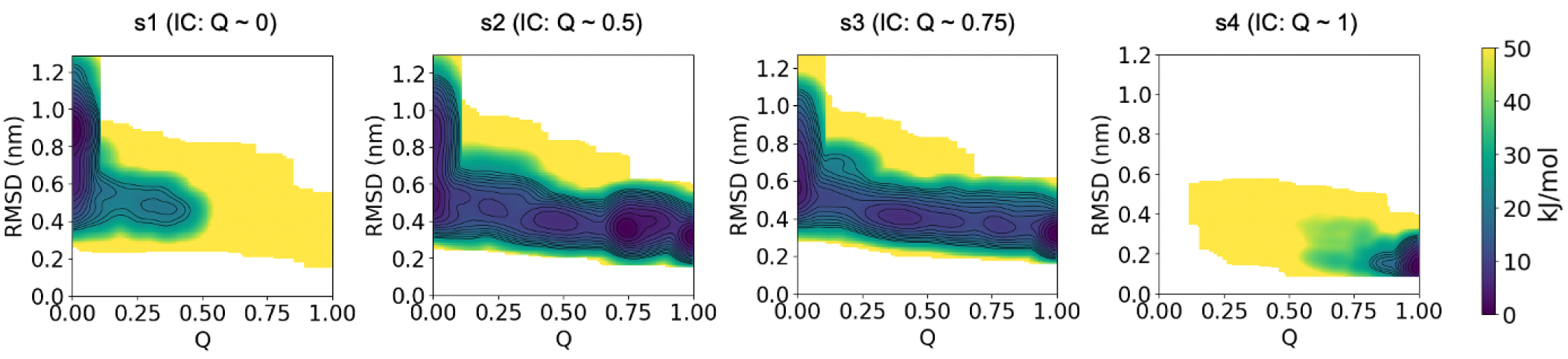
Free energy surfaces (FES) of the GAGA tetraloop as obtained from Deep-DriveMD for the simulation sets s1-s4. Each replica in sets 1, 2, 3, and 4 was simulated for 2.24, 2.25, 1.61, and 2.24 *µ*s, respectively.

**Figure S2:**
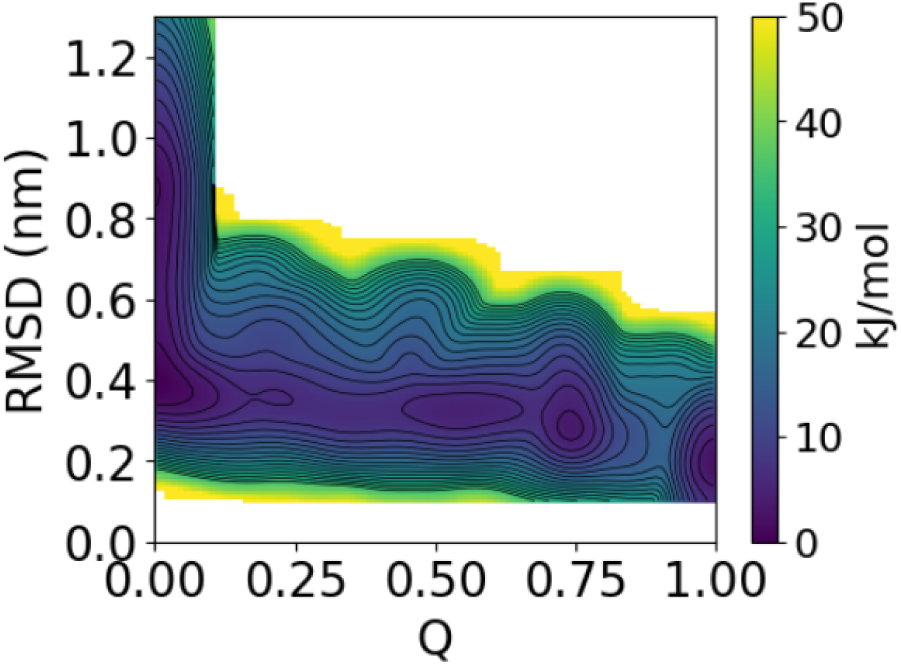
Free energy surface (FES) of the GAGA tetraloop as obtained from DeepDriveMD when the new initial conditions are selected after refining the top outliers with respect to their heavy atom RMSD to the native state, hence leading the simulation towards the folded state.

**Figure S3:**
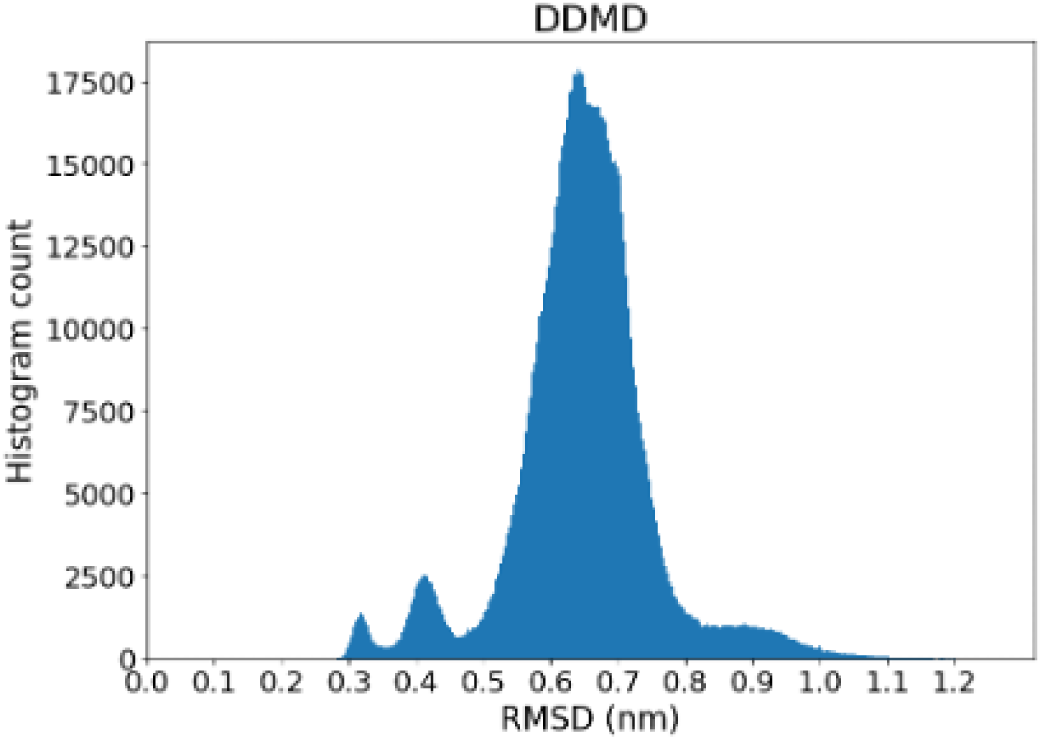
Histograms distribution of the sampled conformations for their heavy atom RMSD to the native state for s3.

**Figure S4:**
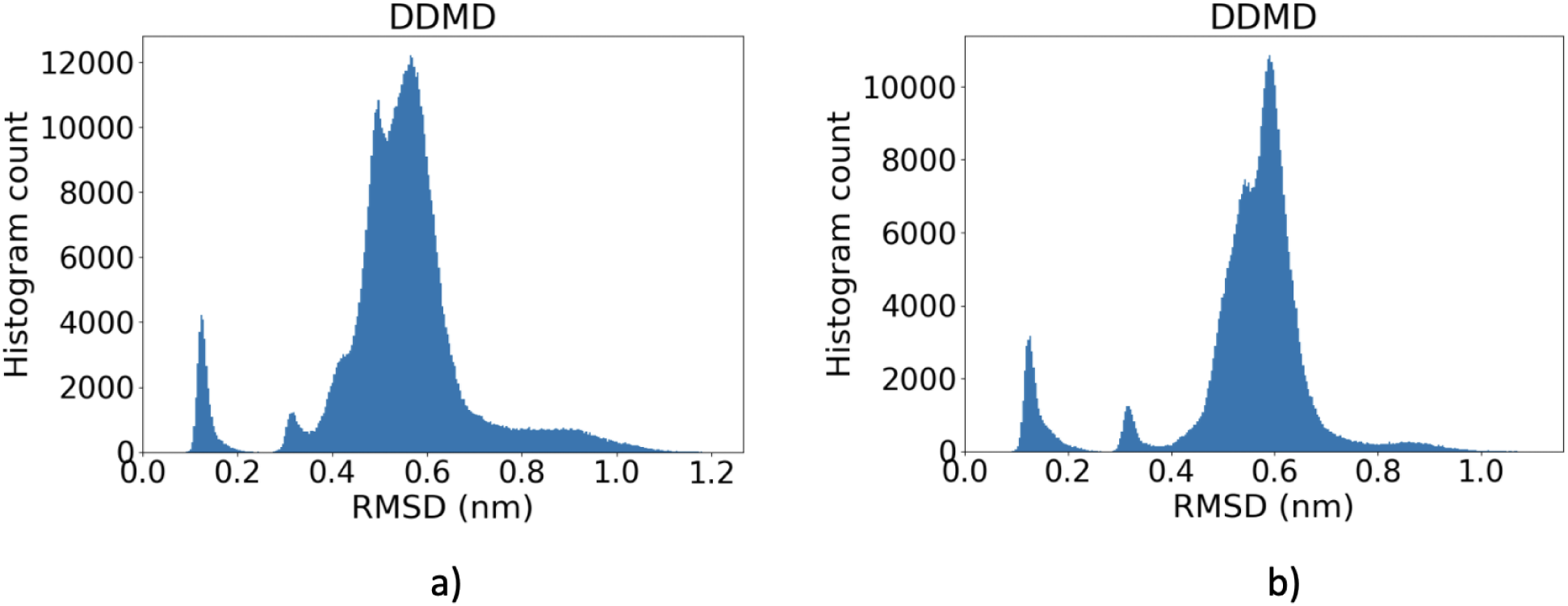
Histograms of the sampled conformations for their heavy atom RMSD to the native state for a) s5 and b) s6.

**Figure S5:**
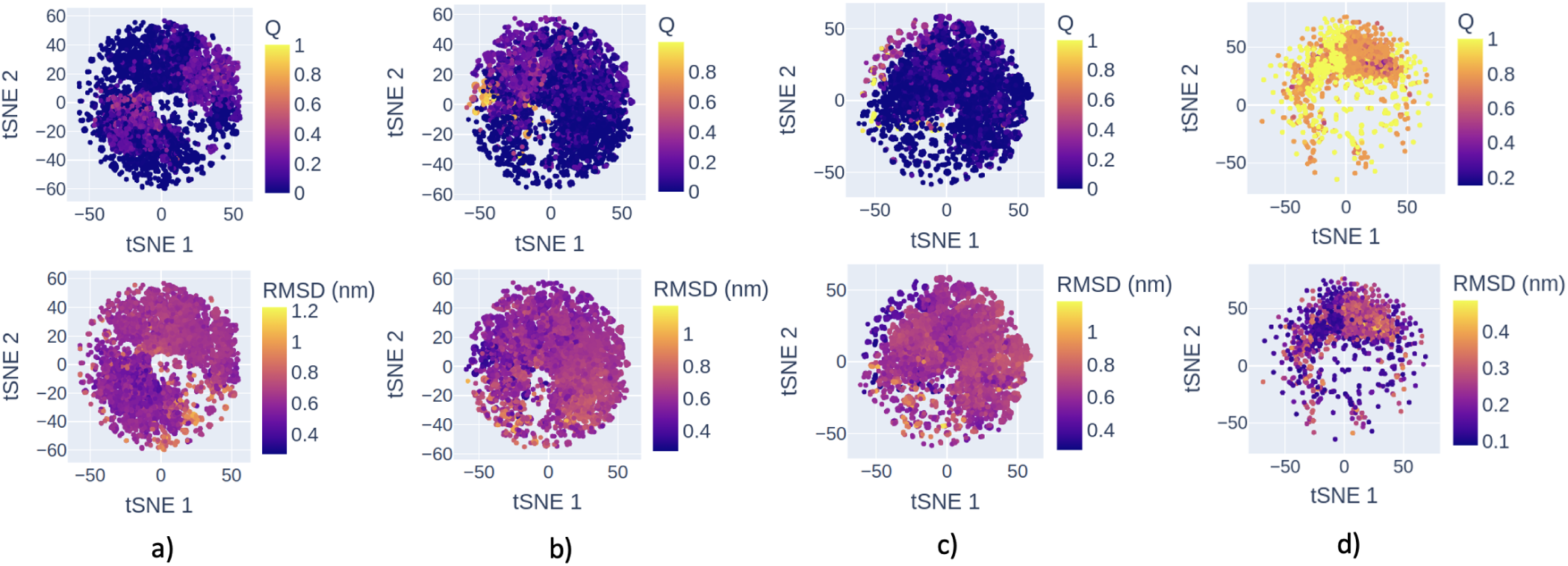
Latent space representations of conformations sampled by a) s1, b) s2, c) s3, and d) s4, projected on a 2-D space using tSNE. The latent space has been colored separately using the two order parameters, Q (top panel) and the heavy atom RMSD to the native state (bottom panel). The plots show that the latent space can segregate the conformations based on the respective order parameter.

**Figure S6:**
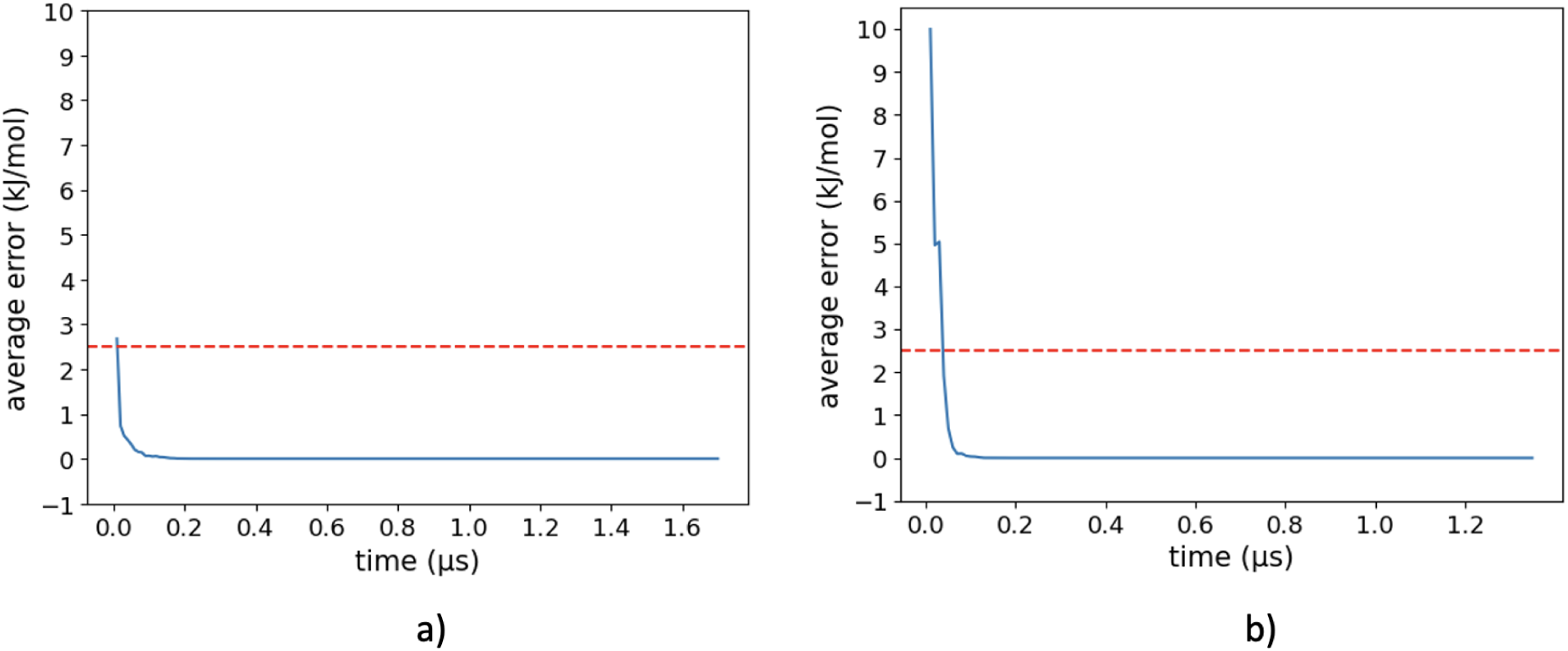
Average free energy error of the FES as a function of time for a) s5, and b) s6, by taking the final FES (i.e. obtained after the total simulation time per replica) as the reference. The average free energy error is capped at 10 kJ/mol. The red line represents the energy of thermal fluctuations (*kT ≈* 2.5 kJ/mol at 300 K). The respective simulation was considered to converge when the average error settled down to *∼* 0 kJ/mol.

## Notes

### Competing Interest Statement

The authors have declared no competing interest.

